# ERGA-BGE genome of *Erebia palarica* Chapman, 1905: a montane butterfly endemic to North-West Iberia

**DOI:** 10.1101/2025.01.08.631636

**Authors:** Marta Vila, Laura Torrado-Blanco, Nils Ryrholm, David Romero-Pedreira, María Conejero, Astrid Böhne, Rita Monteiro, Thomas Marcussen, Rebekah Oomen, Torsten Hugo Struck, Laura Aguilera, Marta Gut, Francisco Câmara Ferreira, Fernando Cruz Rodriguez, Jèssica Gómez-Garrido, Tyler S. Alioto, Chiara Bortoluzzi

## Abstract

The reference genome of *Erebia palarica* will provide valuable insights into evolutionary and conservation genomics. On one hand, the reference genome paves the way to unravel the speciation process, reproductive barriers, and putative hybridisation of *E. palarica* with its closely related sister species: *Erebica meolans*. On the other hand, the reference genome will play an important role in the genetic monitoring of this endemic species, facilitating the use of genomics to estimate population genomics parameters. The genome was assembled into 14 contiguous chromosomal pseudomolecules (Z chromosome included). This chromosome-level assembly encompasses 0.49 Gb, composed of 38 contigs and 17 scaffolds, with contig and scaffold N50 values of 34.2 Mb and 38.4 Mb, respectively.

## Introduction

The butterfly *Erebia palarica* Chapman 1905, also known as Chapman’s ringlet, is endemic to the Cantabrian Mountains (North-West Spain) and surrounding mountain ranges (i.e., Ancares, Courel, Queixa, Montes de León). This univoltine *Satyrinae* species is the largest species of the genus in Europe and it usually occurs at altitudes above 1,000 m. *Erebia palarica* is currently classified as “Least Concern” on the IUCN Red List. However, Torrado-Blanco et al. (2025) suggested the change to “Near Threatened” based on census and contemporary effective population size, as well as other criteria such as predicted reductions of distribution area (Romo et al. 2023) and current area of occupancy and extent of occurrence.

Overall, adults of *E. palarica* flutter in open scrublands, between late June and early September. However, local flight peaks likely extend for a month. Adults are frequently seen near heath (*Erica* L., 1753) and heather (*Calluna* Salisb.) formations, and use rocky trails for basking. This species plays a relevant role in the montane ecosystem, acting as pollinator (adults use nectar resources provided by different plant taxa) and prey species. The larval cycle of *E. palarica* is far from understood (Clarke 2024). Caterpillars have been reported to feed on *Festuca* L., 1753 grasses, but formal confirmation of the mono- or oligophagy is needed (reviewed by Torrado-Blanco 2024). Most populations of *E. palarica* are known to host the same endosymbiont *Wolbachia* Hertig, 1936 strain as the surveyed populations of Iberian *Erebia meolans* (Prunner, 1789), its sister species. The chromosome number is the same for both species (2n = 28; de Lesse 1953, Torrado-Blanco 2024).

The phylogeography of *E. palarica* has been studied with mitochondrial Sanger sequencing and microsatellite markers (Torrado-Blanco et al. 2024). Therefore, developing a high-quality reference genome for *E. palarica* is essential for advancing our understanding of its unique genetic makeup and adaptive traits.

The generation of this reference resource was coordinated by the European Reference Genome Atlas (ERGA) initiative’s Biodiversity Genomics Europe (BGE) project, supporting ERGA’s aims of promoting transnational cooperation to promote advances in the application of genomics technologies to protect and restore biodiversity (Mazzoni et al. 2023).

## Materials & Methods

ERGA’s sequencing strategy includes Oxford Nanopore Technology (ONT) and/or Pacific Biosciences (PacBio) for long-read sequencing, along with Hi-C sequencing for chromosomal architecture, Illumina Paired-End (PE) for polishing (i.e. recommended for ONT-only assemblies), and RNA sequencing for transcriptome profiling, to facilitate genome assembly and annotation.

### Sample and Sampling Information

On 2 July 2022, four female specimens of *E. palarica* were sampled by Marta Vila and David Romero-Pedreira at Serra do Courel, Lugo, Spain: three from A Cabeza Grande and one from Alto do Couto. On 20 July 2023, a fifth female was sampled by Marta Vila and Laura Torrado-Blanco at A Cabeza Grande. The identification of the species was confirmed based on morphology by Marta Vila, Laura Torrado-Blanco and Nils Ryrholm, and barcoding by María Conejero.

The sampling of all five female specimens was conducted under permits LU/003/2022 and LU/012/2023, issued by the Xunta de Galicia, Consellería de Medio Ambiente, Territorio e Vivenda, Servizo de Patrimonio Natural (Spain). Sampling was performed using a butterfly net. Specimens were handled in two ways before storage at -80°C. Two females collected at A Cabeza Grande in 2022 were immediately euthanized using dry ice, kept there for 34 hours, and then transferred to -80°C. The remaining three specimens were maintained alive for 24 hours until arrival at the lab and then frozen at -80°C. All samples were stored at -80°C until DNA extraction.

#### Vouchering information

Physical reference materials for a proxy male (sampled under the aforementioned permits) specimen have been deposited in the Museo Nacional de Ciencias Naturales (MNCN, CSIC) https://mncn.csic.es/en under the accession number MNCN:Ent:366619.

#### Frozen reference tissue material

The whole organism is available in the Museo Nacional de Ciencias Naturales (MNCN, CSIC) https://mncn.csic.es/en under tissue voucher ID MNCN:Ent:366618 and proxy tissue voucher ID MNCN:ADN:151.720.

### Data Availability

*Erebia palarica* and the related genomic study were assigned to Tree of Life ID (ToLID) ‘ilErePala4’ and all sample, sequence, and assembly information are available under the umbrella BioProject PRJEB75270. The sample information is available at the following BioSample accession: SAMEA114504880. The genome assembly is accessible from ENA under accession number GCA_964035005.2 and the annotated genome is available through the Ensembl Beta page (https://beta.ensembl.org/). Sequencing data produced as part of this project are available from ENA at the following accessions: ERX12324509, ERX12324510, ERX12324511, ERX13168336, and ERX13168337. Documentation related to the genome assembly and curation can be found in the ERGA Assembly Report (EAR) document available at https://github.com/ERGA-consortium/EARs/blob/main/Assembly_Reports/Erebia_palarica/ilErePala4/ilErePala4_EAR.pdf. Further details and data about the project are hosted on the ERGA portal at https://portal.erga-biodiversity.eu/organism/SAMEA114504880.

### Genetic Information

The estimated genome size, based on ancestral taxa, is 0.48 Gb. This is a diploid genome with a haploid number of 14 chromosomes (2n = 28) and sex chromosomes (W and Z) based on previous studies on closely related species (Lohse et al. 2022, Bioproject PRJEB80714). All information for this species was retrieved from Genomes on a Tree (Challis et al. 2023).

### DNA/RNA processing

DNA was extracted from head and thorax using the Blood & Cell Culture DNA Mini Kit (Qiagen) following the manufacturer’s instructions. DNA quantification was performed using a Qubit dsDNA BR Assay Kit (Thermo Fisher Scientific) and DNA integrity was assessed using a Femtopulse system (Genomic DNA 165 Kb Kit, Agilent). DNA was stored at 4ºC until use.

RNA was extracted using an RNeasyMini Kit (Qiagen) according to the manufacturer’s instructions. RNA was extracted from three different tissues: leg, wing and abdomen. RNA quantification was performed using the Qubit RNA BR Kit and RNA integrity was assessed using a Fragment Analyzer system (RNA 15nt Kit, Agilent). RNA was pooled in a 1:1:2 (leg:wing:abdomen) ratio before library preparation and stored at -80ºC until use.

### Library Preparation and Sequencing

For long-read whole genome sequencing, a library was prepared using the SQK-LSK114 kit (Oxford Nanopore Technologies, ONT) and it was sequenced on a PromethION 24 A series instrument (ONT). A short-read WGS library was prepared using the KAPA Hyper Prep Kit (Roche). The RNA library from the pooled sample was prepared using the KAPA mRNA Hyper prep kit (Roche). A Hi-C library was prepared from abdomen tissue using the ARIMA High Coverage Hi-C Kit (ARIMA), followed by the KAPA Hyper Prep Kit for Illumina sequencing (Roche). All the short-read libraries were sequenced on a NovaSeq 6000 instrument (Illumina).

In total 104X Oxford Nanopore, 64X Illumina WGS shotgun, and 81X HiC data were sequenced to generate the assembly.

### Genome Assembly Methods

The genome was assembled using the CNAG CLAWS pipeline (Gomez-Garrido 2024). Briefly, reads were preprocessed for quality and length using Trim Galore v0.6.7 and Filtlong v0.2.1, and initial contigs were assembled using NextDenovo v2.5.0, followed by polishing of the assembled contigs using HyPo v1.0.3, removal of retained haplotigs using purge-dups v1.2.6 and scaffolding with YaHS v1.2a. Finally, assembled scaffolds were curated via manual inspection using Pretext v0.2.5 with the Rapid Curation Toolkit (https://gitlab.com/wtsi-grit/rapid-curation) to remove any false joins and incorporate any sequences not automatically scaffolded into their respective locations in the chromosomal pseudomolecules (or super-scaffolds). Finally, the mitochondrial genome was assembled as a single circular contig of 15,221 bp using the FOAM pipeline (https://github.com/cnag-aat/FOAM) and included in the released assembly (GCA_964035005.2). Summary analysis of the released assembly was performed using the ERGA-BGE Genome Report ASM Galaxy workflow (https://doi.org/10.48546/WORKFLOWHUB.WORKFLOW.1103.2).

## Results

### Genome Assembly

The genome assembly has a total length of 493,162,700 bp in 17 scaffolds (Z chromosome included) and the mitogenome (Figures 1 & 2), with a GC content of 36.9%. The assembly has a contig N50 of 34,188,440 bp and L50 of 6 and a scaffold N50 of 38,441,827 bp and L50 of 6. The assembly has a total of 21 gaps, totaling 4.2 kb in cumulative size. The single-copy gene content analysis using the Lepidoptera database with BUSCO v5.5.0 (Manni et al. 2021) resulted in 98.3% completeness (97.8% single and 0.5% duplicated). 87.7% of reads k-mers were present in the assembly and the assembly has a base accuracy Quality Value (QV) of 43.9 as calculated by Merqury (Rhie et al. 2020).

**Figure 1.**
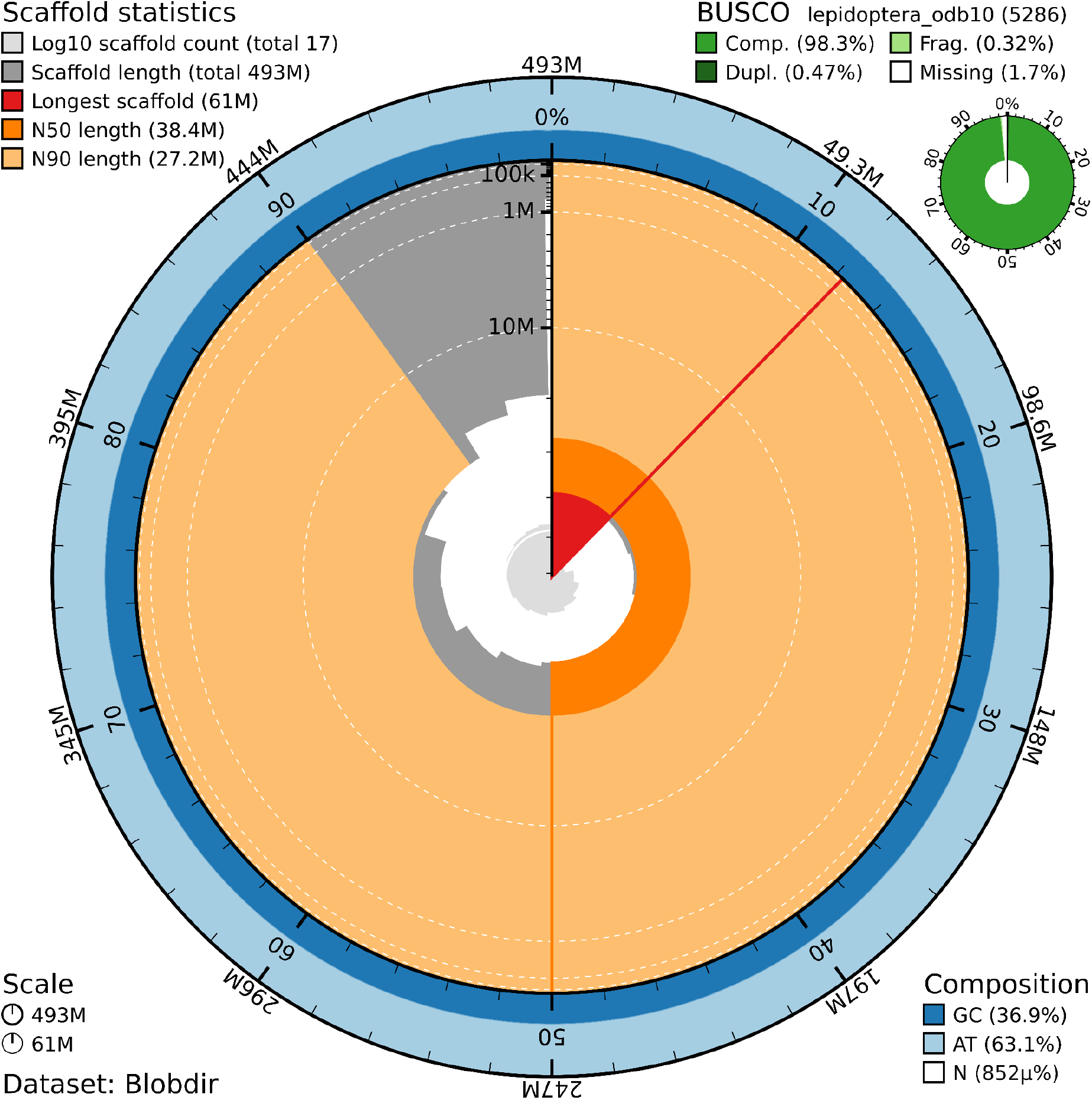
Snail plot summary of assembly statistics. The main plot is divided into 1,000 size-ordered bins around the circumference, with each bin representing 0.1% of the 493,162,700 bp assembly including the mitochondrial genome. The distribution of sequence lengths is shown in dark grey, with the plot radius scaled to the longest sequence present in the assembly (61.0 Mb shown in red). Orange and pale-orange arcs show the scaffold N50 and N90 sequence lengths (38,441,827 and 27,179,420 bp), respectively. The pale grey spiral shows the cumulative sequence count on a log-scale, with white scale lines showing successive orders of magnitude. The blue and pale-blue area around the outside of the plot shows the distribution of GC, AT, and N percentages in the same bins as the inner plot. A summary of complete, fragmented, duplicated, and missing BUSCO genes found in the assembled genome from the Lepidoptera database (odb10) is shown in the top right.

**Figure 2.**
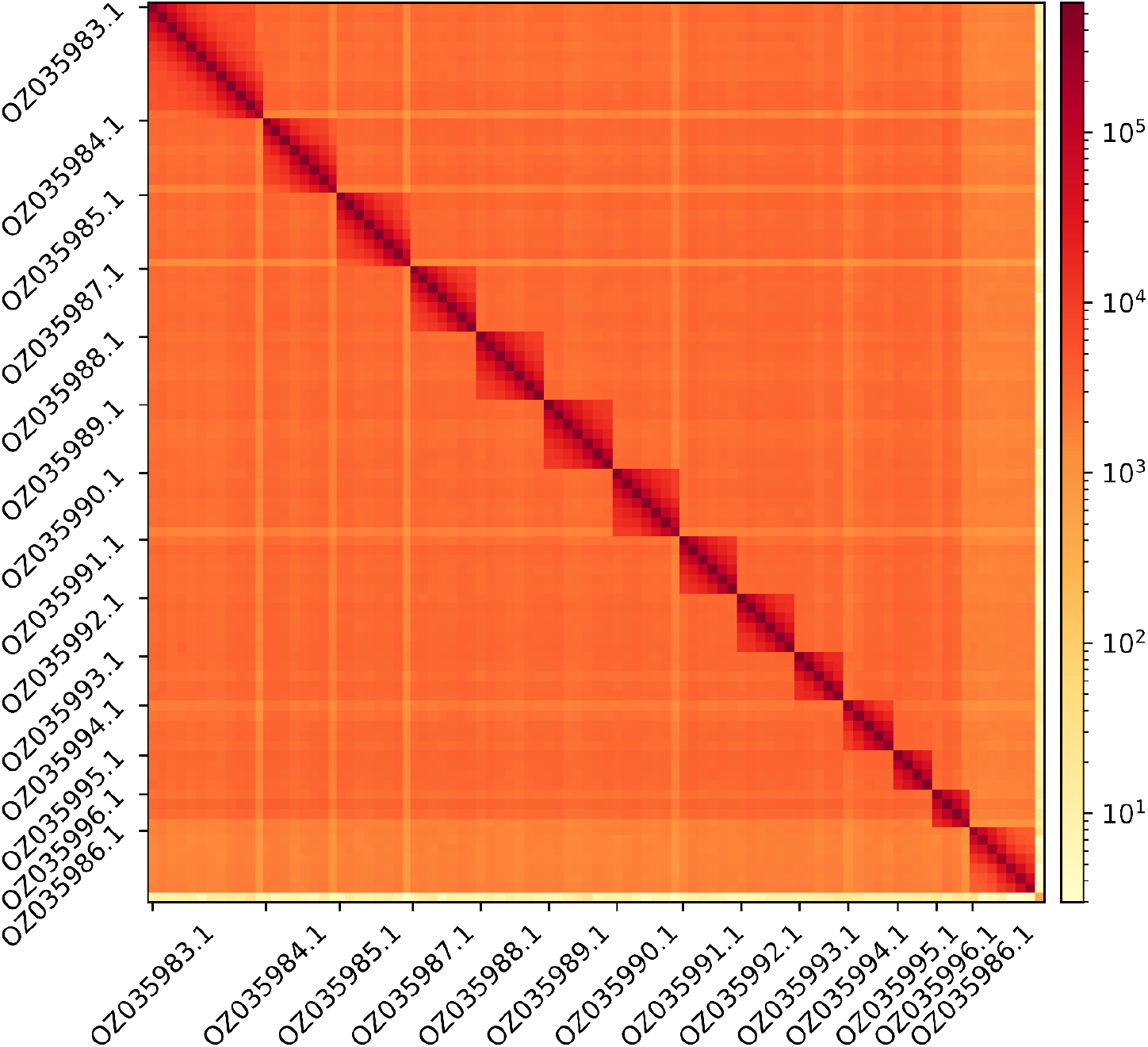
Hi-C contact map showing spatial interactions between regions of the genome. The diagonal corresponds to intra-chromosomal contacts, depicting chromosome boundaries. The frequency of contacts is shown on a logarithmic heatmap scale. Hi-C matrix bins were merged into a 100 kb bin size for plotting. The Hi-C contact map shows the GenBank names of the 13^th^ largest chromosomes and the Z chromosome (GenBank name: OZ035986.1).

## Acknowledgements

We would like to express our gratitude to Dr. Rodolfo Barreiro and Dr. Mercedes París for their assistance with sample storage and vouchering, respectively. We appreciate the suggestions from Nadir Alvarez and ERGA-Spain. We would like to acknowledge the assembly reviewer, Thomas Brown, from the Leibniz Institute for Zoo and Wildlife Research (Germany). The authors acknowledge the support of the Freiburg Galaxy Team: Saim Momin and Björn Grüning, Bioinformatics, University of Freiburg (Germany), funded by the German Federal Ministry of Education and Research BMBF grant 031 A538A de.NBI-RBC and the Ministry of Science, Research and the Arts Baden-Württemberg (MWK) within the framework of LIBIS/de.NBI Freiburg.

## Author Contributions

MV coordinated the project; MV, LT-B, and DR-P collected the species; MV, LT-B, and NR identified the species; MC did the barcoding; MV and LT-B sampled and preserved biological material and provided metadata; RM, TM, RO, THS, and AsB provided support in sampling, shipping of biological material, metadata collection, and management; LA and MG extracted DNA, prepared libraries, and performed sequencing; FC, FCF and JGG performed genome assembly and curation under the supervision of TSA; CB generated the analysis and report. All authors contributed to the writing, review, and editing of this genome note and read and approved the final version.

